# Altered Cortical Network Dynamics during Observing and Preparing Action in Patients with Corticobasal Syndrome

**DOI:** 10.1101/2024.07.19.604284

**Authors:** Marius Krösche, Christian Hartmann, Markus Butz, Alfons Schnitzler, Jan Hirschmann

## Abstract

Besides parkinsonism, higher order cortical dysfunctions such as apraxia are hallmarks of the corticobasal syndrome (CBS). To date, little is known about the electrophysiological underpinnings of these symptoms.

To shed more light on the pathophysiology of CBS, we recorded the magnetoencephalogram of 17 CBS patients and 20 age-matched controls engaged in an observe-to-imitate task. The task involved the display of a tool-use video in first person view (action observation), a written instruction to withhold movement until the presentation of a Go cue (movement preparation), and unilateral tool-use imitation. We investigated modulations of spectral power on the source level.

Action observation was associated with an event-related desynchronization in the beta-band (13-30Hz), which was weaker in CBS patients than in healthy controls. The group effect localized to superior parietal, primary motor, premotor and inferior frontal cortex bilaterally. While participants awaited the Go cue, beta power was again suppressed in the hemisphere contralateral to movement, and the rate of suppression correlated with reaction time. This modulation, too, was weaker in the CBS group. Immediately before movement onset, however, beta power was similar in both groups.

Our results reveal that action observation triggers beta suppression, likely reflecting motor cortical disinhibition, which is reduced in CBS patients. This pathological alteration might be a neural correlate of a selective deficit in the embedding of observed action into a motor representation (visuo-motor mapping). The suboptimal timing of the beta-power suppression to the upcoming Go cue presumably reflects a deficit in learning the trial’s temporal structure rather than a deficit in movement initialization.

**Highlights:**

- Action observation elicits a suppression of cortical beta power.
- This suppression is diminished in CBS patients relative to controls.
- The timing of beta desynchronization to an upcoming Go cue is impaired in CBS patients.
- No evidence for an alteration of movement initialization in CBS patients.

## Introduction

The corticobasal syndrome (CBS) is a rare neurodegenerative disease that rapidly progresses, with no causal treatment options available to this day (Armstrong et al. 2013). The cardinal clinical features are parkinsonism, cognitive decline, and apraxia, i.e. the inability to enact skilled movements despite intact primary sensory and motor function (Park 2017), often asymmetrically presented at disease onset (Armstrong et al. 2013). At first clinical presentation, CBS is frequently misdiagnosed due to its variable symptomology (Osaki et al. 2004; Joutsa et al. 2014; Alexander et al. 2014; Aiba et al. 2023). The cardinal neuropathological hallmark is the accumulation of misfolded tau-protein in neurons and glia cells, followed by their degeneration (Höglinger et al. 2018). The protein-pathology presumably begins subcortically in the basal ganglia and spreads to various cortical sites, with widespread effects on the frontal, parietal and temporal lobes (Leuzy et al. 2019). Especially motor areas like peri-rolandic, premotor and supplementary motor regions are frequently mentioned as cortical hubs of tau pathology (Pardini et al. 2019; Cho et al. 2017; Kikuchi et al. 2016; Smith et al. 2017) and degeneration (Huey et al. 2009; Josephs et al. 2010; Whitwell et al. 2010; Dutt et al. 2016; Matsuda et al. 2020).

The electrophysiology of CBS is not well investigated. Electroencephalography (EEG) and magnetoencephalography (MEG) studies hint towards widespread spectral slowing of brain activity at rest (Tashiro et al. 2006; Barcelon et al. 2019; Krösche et al. 2023), pronounced in frontal and parietal sites (Krösche et al. 2023). The clinical consequences of these alterations, however, remain unclear. Among various cognitive functions, frontoparietal networks are implicated in the processes of action observation (Molenberghs et al. 2012; Hardwick et al. 2018), motor imagery (Caspers et al. 2010; Hétu et al. 2013; Hardwick et al. 2018) and action execution (Jeannerod 2001; Hardwick et al. 2018). All of these processes are associated with a desynchronization of motor rhythms in the alpha and beta range (Schnitzler et al. 1997; McFarland et al. 2000; Caetano et al. 2007; Fairhall et al. 2007; Eaves et al. 2016). On a cellular level, these functions are presumably supported by mirror neurons, i.e. neurons that discharge similarly during execution and observation of goal-directed movements (Di Pellegrino et al. 1992; Gallese et al. 1996; Rizzolatti et al. 1996). Mirror neurons were first described in nonhuman primates in frontal and parietal regions (Di Pellegrino et al. 1992; Gallese et al. 1996; Fogassi et al. 2005) before their discovery in humans (Mukamel et al. 2010). Their activity represents the observed action in a motoric neuronal code (Rizzolatti et al. 1996; Rizzolatti et al. 2009; Heyes and Catmur 2022). Damage to the mirror neuron system may lead to deficits in performing and perceiving goal-directed movements in CBS.

In line with this concept, neuropathology studies demonstrated that cortical degeneration in frontoparietal areas is related to apraxia (Gross and Grossman 2008; Park 2017). The same areas show disease-related structural changes in CBS (Huey et al. 2009). On the electrophysiological level, a previous study found pathologically increased left parietal to right premotor beta-band coherence (13-30Hz) prior to tool-use pantomime in three CBS patients with apraxia (Wheaton et al. 2008). While these observations align well with the established role of frontoparietal networks in goal-directed movement, there is not enough data available to draw firm conclusions on how activity in these networks relates to CBS symptoms.

To help fill this knowledge gap, the current study investigated the association between oscillatory activity and deficits in action observation and movement preparation in a comparably large sample of CBS patients. We made use of an observe-to-imitate task, engaging frontoparietal networks presumed to be dysfunctional in CBS patients.

## Methods

### Participants

In total, 17 CBS patients and 20 healthy controls performed the imitation task. Data from two control participants were discarded. One participant was taking antidepressants and another one had aphantasia i.e. was impaired in motor imagery (Dupont et al. 2022). Four patients of the CBS group were excluded. Two patients did not follow the task instructions and two further patients were diagnosed with Progressive Supranuclear Palsy or Multisystem Atrophy later during clinical follow-up. Consequently, the data of 13 CBS patients and 18 control subjects were used for analysis. The groups did not differ in age (see Table 1.; *t(29) = -1.628, p = 0.114*). In the patient group, we performed several neuropsychological / neurological tests to evaluate cognitive impairment, parkinsonism, and apraxia.

**Table 1:** Summary data of the study cohorts.

| Group | Mean age (years) | Gender (f/m) | Diagnosis (poss./prob.) | Mean (Std.) UPDRS-III (sum) | Mean (Std.) MoCA (normalized) | Mean (Std.) Goldenberg (sum) | Mean (Std.) TULIA (sum) |
| --- | --- | --- | --- | --- | --- | --- | --- |
| CBS | 65.23 | 6/7 | 5/8 | 35.92 (22.03) | 0.66 (0.21) | 56 (21.73) | 18.25 (5.86) |
| HC | 69.17 | 10/8 | - | - | - | - | - |

CBS: Corticobasal Syndrome, HC: Healthy controls.

We used the Montreal Cognitive Assessment (MoCA) to evaluate cognitive impairment (Nasreddine et al. 2005), the UPDRS-III for the severity of parkinsonism (Goetz et al. 2008), the Goldenberg’s Apraxia Test (Goldenberg 1996) and the Test of Upper Limb Apraxia (Vanbellingen et al. 2010) for apraxia. We report relative test scores for cognitive impairment as physical disabilities prevented testing items on visuospatial orientation reliably in two patients. The MoCA scores were normalized by dividing the achieved score by the maximal score, considering only scorable items. The local ethics committee approved the study (study-number: CBS: 2019-447-andere) and every participant gave written informed consent prior to participation, in accordance with the Declaration of Helsinki.

### Trial Design

We made use of an observe-to-imitate task, i.e. the observation of an action, followed by a delayed request to imitate that same action. Each trial began with the display of a fixation cross (variable stimulus duration: 1-3s) followed by a 2s video displaying a person using a hammer or a screwdriver, either with the left or with the right hand, in first-person view. This was followed by text instructing the participants to withhold movement *(“do not move yet”*) until a Go cue appeared (movement preparation phase, duration: 5s). The Go cue was on screen for 4s. Meanwhile, participants imitated the action with the cued hand until a Stop cue was presented (stimulus duration: 1s).

Following 10 practice trials, maximally 4 blocks of 40 trials (21 screwdriver, 19 hammer trials in random order) were recorded per participant. In each block, participants were requested to respond with the right hand or with the left hand only (first block randomized, hand switch after each subsequent block). We recorded at least one right hand block and one left hand block in all but one patient, who completed only one left hand block (median: 4 blocks; Range: 1 – 4; Supplementary Table 1). All healthy controls completed all four blocks.

### Recordings

Brain activity was recorded with a 306-sensor MEG system (VectorView, MEGIN, Espoo, Finland) in a magnetically shielded room, with a sampling rate of 1000Hz. During the recordings, participants were sitting in upright position with their arms laying on a table in front of them. Electromyograms were recorded from both forearms and accelerometers were attached to the left and right index finger to track upper limb motion. In addition, we recorded a vertical and a horizontal electrooculogram.

### Data Preprocessing

#### Data cleaning

Data analysis was performed with MATLAB 2018a (MathWorks, Natick, MA), Python 3.9.1, and the Fieldtrip toolbox (version 18.01.2023, Oostenveld et al. 2011). After discarding bad channels, we applied temporal Signal Space Separation (*tSSS*, Taulu and Simola 2006) to attenuate the interference of sources from outside the MEG helmet (MNE-Python Toolbox 1.3.1, Gramfort et al. 2013). Next, we filtered the *tSSS*-cleaned data with the spectral interpolation algorithm (Leske and Dalal 2019) to remove line noise and its harmonics and applied a high-pass filter with a cut-off frequency of 0.5Hz before resampling the data to 250Hz. Next, we screened the data, removed periods of movement artifacts and sensor noise, and applied independent component analysis. Independent components of non-brain origin, e.g. heartbeat and eye movements, were discarded (median number of ICs removed: 2.5, range: 1 – 5).

#### Trial definition

We restricted data analysis to the pre-movement period to avoid movement artifacts. The preprocessed data were segmented into trials of 14s, centered on the Go cue, encompassing baseline (-7.75s to -7s), video (-7s to -5s), and movement preparation phase (-5s to 0s).

We screened all trials and discarded trials containing movement artifacts, particularly when artifacts occurred within the baseline period. To detect outliers in the baseline period, we applied a semiautomatic procedure involving a threshold applied to the first principal component of the smoothed accelerometer signals (+/-2.33 SD). If ≥100ms of data within or prior to the baseline period (-9s to - 7s) were marked as outliers the trial was considered invalid. The results were visually checked and corrected if necessary. In total, 22.89% of trials were removed (CBS: 27.27%, HC: 20.43%). This meticulous screening procedure served to ensure that the data analyzed here do not contain movement artifacts.

#### Movement onset detection

The Teager Kaiser Energy Operator (TKEO, Solnik et al. 2010) was computed to determine movement onset in the accelerometer signals. TKEO was z-normalized, using the mean of a movement-free reference period (-3 to -1 s with respect to the Go cue), and a threshold was applied to determine movement onset (>200 SD TKEO of non-moving hand). Movement onset estimates were visually inspected and corrected if necessary.

### Source Reconstruction

T1-weighted magnetic resonance imaging (MRI) scans (Siemens Magnetom Tim Trio, 3-T MRI scanner, Munich, Germany) were used to compute individualized, single-shell head models (Nolte 2003). Individual MR-images were available for all CBS patients and for 15 of 19 healthy controls. For the remaining participants, we used a template brain (Holmes et al. 1998). The coordinate systems of the MEG and the MRI data were aligned based on anatomical landmarks sampled with a digitizer system (Isotrak, Polhemus, Colchester, Vermont, USA) prior to the MEG recordings. Trials were cut into nonoverlapping segments of 1s length, and we computed one covariance matrix per experimental block. Based on the gradiometer covariance, we estimated source activity for 567 positions on the cortical surface in Montreal Neurological Institute (MNI) space, using a Linearly Constrained Minimal Variance beamformer and a regularization parameter of 5% (van Veen et al. 1997). Subsequently, we applied singular value decomposition to the x-, y-, and z-components of the resulting dipoles and kept the vector with the largest eigenvalue.

### Spectral Analysis

Time-frequency decomposition of the source-reconstructed trial data was done via Morlet wavelets for frequencies ranging between 4Hz and 90Hz, in 1Hz steps. The number of cycles for the wavelets ranged from 4 to 15, increasing as a function of frequency in logarithmic space. Wavelets were shifted in steps of 32ms with respect to the signal.

### Statistical Analysis

We applied two-tailed cluster-based permutation tests with Monte-Carlo sampling to quantify between-group effects (Maris and Oostenveld 2007). In short, the dependent variable was shuffled across groups ≥ 50000 times. Each time, a cluster-forming threshold was applied, *t*-values were summed for each cluster and the largest sum was kept, contributing to the empirical null distribution. Clusters in the original (non-shuffled) data were considered significant if their cluster sum fell within the extreme 5% of the null distribution.

As we did not observe differences between hammer and screwdriver trials, we pooled trials across tools for statistical analysis. Similarly, we pooled right hand and left-hand trials after mirroring the activity recorded in left hand trials across the midsagittal plane . Both steps served to increase the signal-to-noise ratio.

#### Estimating the rate of beta power suppression

We used linear regression to estimate the rate of beta event-related desynchronization (ERD) per participant. Before slope estimation, the beta ERD was smoothed with a moving average filter to denoise the signal (width: 160ms). Slope estimates were compared between groups with independent sample *t*-tests at an alpha-error rate of 5%. When relating the beta slopes to reaction time, we pooled values across groups and computed Pearson partial correlation coefficients to control for possible group differences in slope.

#### Bayesian statistics

We applied a Bayesian Analysis of Variance to assess the presence vs. absence of group effects in different phases of the trial. The analysis was performed with R (version 4.4.0) using the BayesFactor package (version 0.9.12-4.7). The model included *group* (HC vs. CBS), *hemisphere* (contralateral vs. ipsilateral) and *trial phase* (action observation, motor preparation, movement initialization) as fixed effects, alongside *participant* (participant ID) as a random effect to account for within-subject variability. The reported Bayes Factors (BF) quantify the evidence for (H1) or against a group difference (H0) per trial phase. They result from post-hoc pairwise comparisons, conducted with Bayesian, independent sample t-tests, after confirming a *group* x *trial phase* interaction. The reporting of evidence follows the recommendations of Andraszewicz et al. (2015).

## Results

### Behavioral Results

All patients were able to imitate hammer and screwdriver use, with large variability in the quality of imitation. Rather than addressing imitation per se, we focused on pre-movement brain activity and behavior.

In the pre-movement phase, both groups revealed slight hand motion in some of the trials, despite being instructed to withhold movement until Go cue presentation. This motion was successfully removed in data cleaning (Fig. 2). After data cleaning, the accelerometer data were largely similar for CBS and HC except for the period immediately preceding the Go cue (cluster: *tsum = -527.109, p = 0.005*; Fig. 2A). Around this time, controls, but not CBS patients, had already initiated their response. Accordingly, controls reacted faster to the Go cue than CBS patients (*t(29) = 4.124, p < 0. 001*, Fig. 2B).

**Figure 1.**
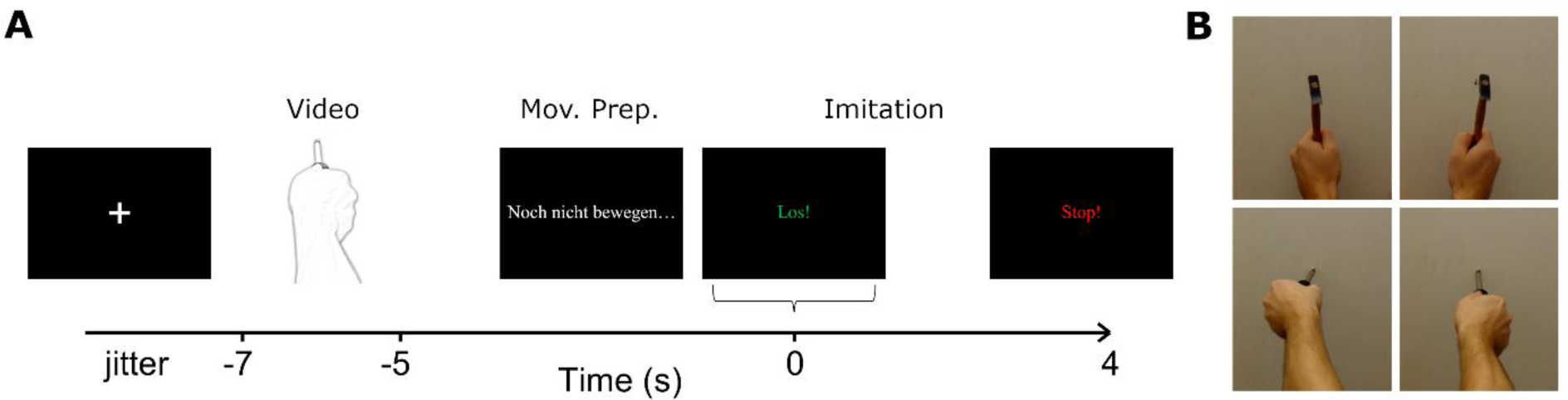
Trial design. A) Following the display of the fixation cross (1-3s), a 2s tool-use video was displayed involving either a hammer or a screwdriver, operated with the left or right hand, depending on the experimental block. This was followed by the instruction to withhold movement (“Noch nicht bewegen…”; 5s) and a Go cue (“Los!”; 4s). Patients were to imitate the action displayed earlier until presentation of a Stop cue (1s). A trial lasted between 13s and 15s. B) A screenshot of each of the four videos used in the task. Mov. Prep.: Movement preparation.

**Figure 2.**
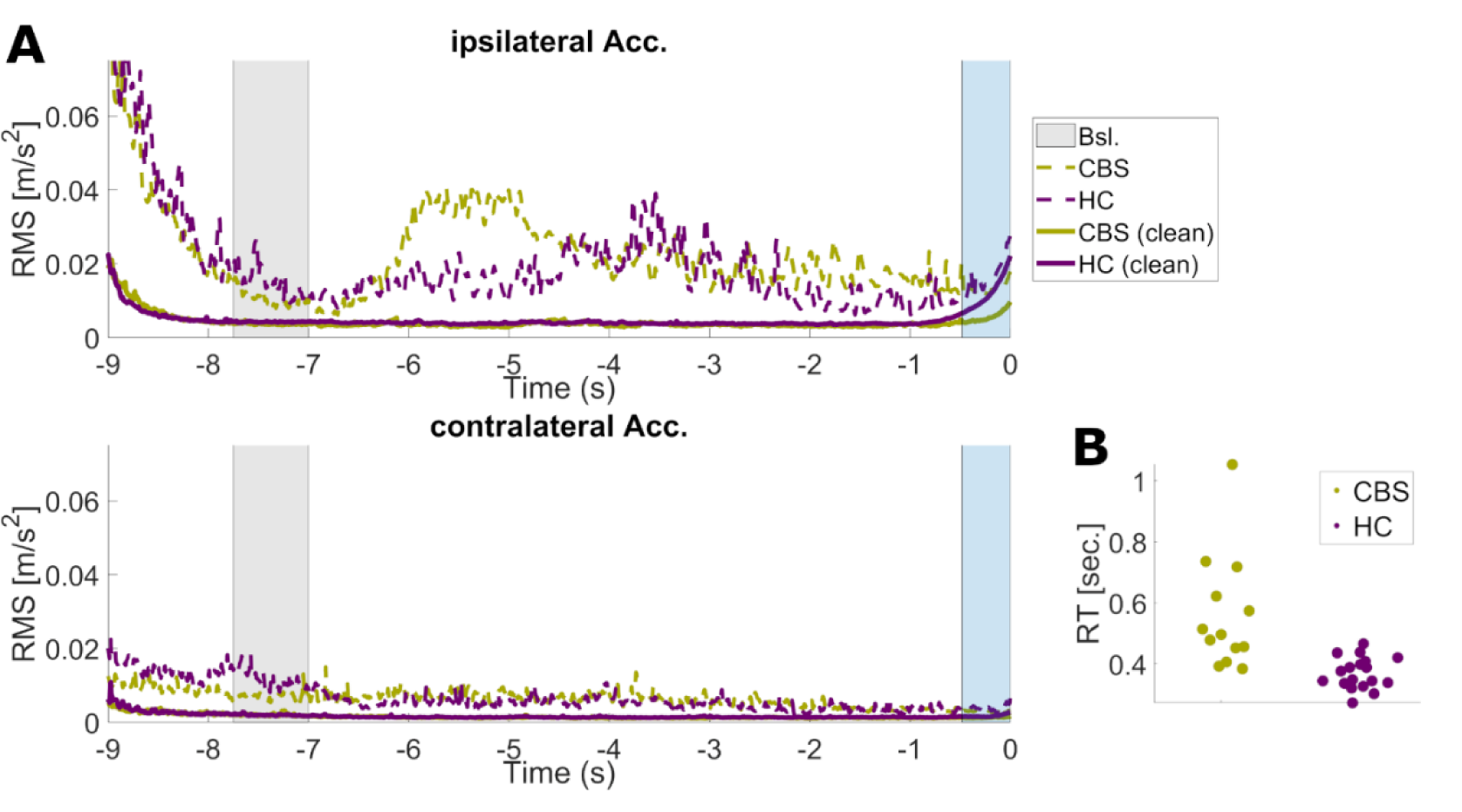
CBS patients reacted slower than healthy controls (HC). A) Root mean square (RMS) accelerometer data for the moving (ipsilateral) and the non-moving (contralateral) hand before (dashed lines) and after (solid lines) cleaning. HC presented larger RMS values than patients within the [-0.48s to 0s] interval (blue shading), with 0s indicating Go cue onset. B) Distribution of trial median reaction time per participant for CBS (median: 0.496s, range: 0.384s to 1.054s) and HC (median: 0.362s, range: 0.272s to 0.466s). Bsl.: Baseline; CBS: Corticobasal Syndrome; HC: Healthy controls; RT: Reaction time.

### Action observation

Action observation was associated with a decrease in alpha/beta power (9-30Hz), henceforth referred to as event-related desynchronization (ERD). The ERD occurred predominantly in sensorimotor and parietal areas, and, with lower amplitude, in temporal and occipital areas (Fig. 3, left panel). It was bilaterally distributed, with a slight emphasis on the hemisphere contralateral to movement, particularly in controls. The desynchronization outlasted the video by ∼800ms in both groups (Fig. 3).

**Figure 3.**
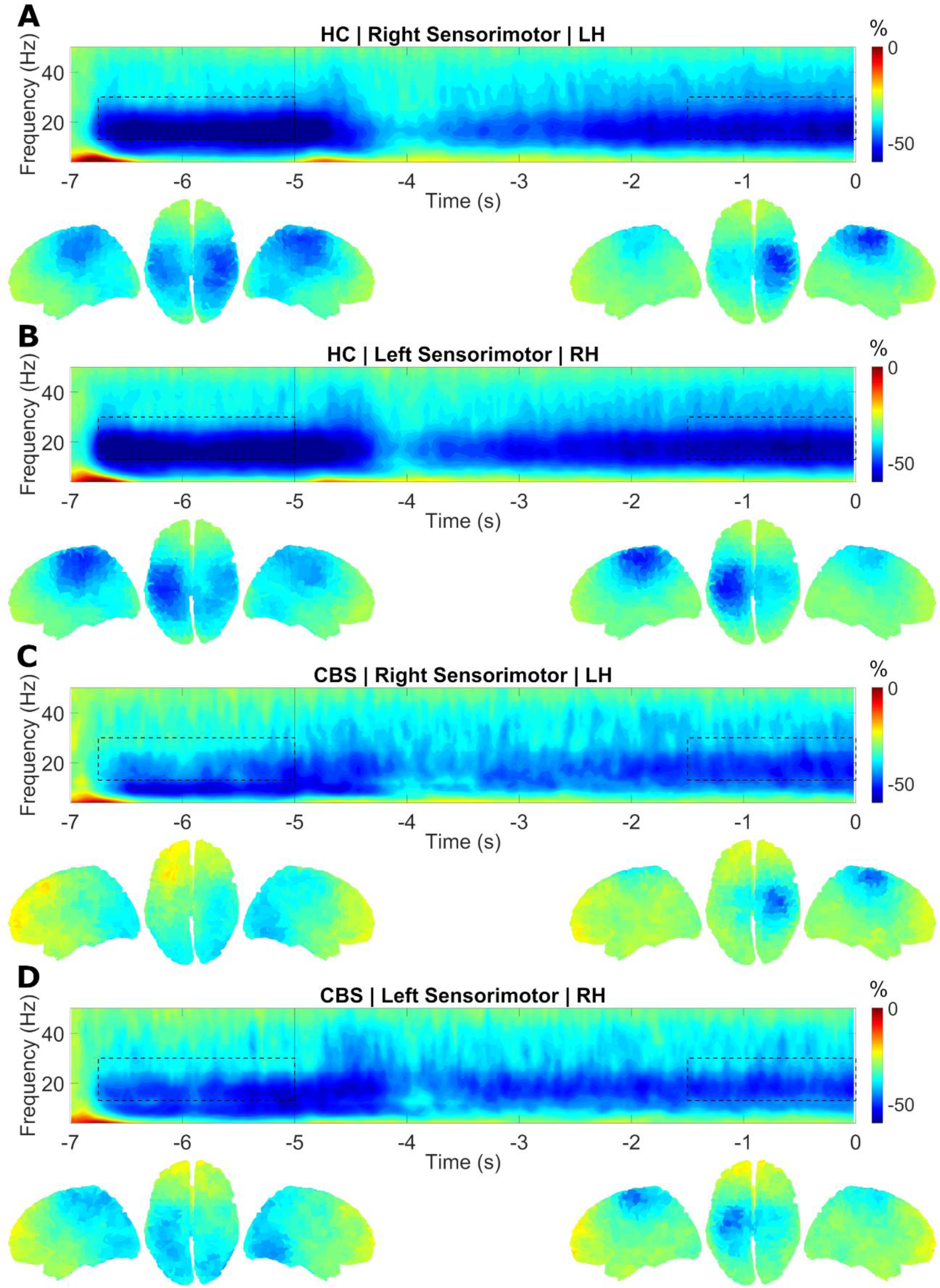
**Action observation and movement preparation were associated with a decrease of beta power in sensorimotor cortex**. Each panels depicts the group-average time-frequency spectrum of sensorimotor cortex contralateral to imitation bodyside. Relative power change with respect to baseline (-7.75s to -7s) is color-coded. The dashed boxes indicate the timefrequency selection used in the source plots below. Participants had to contribute at least 25 trials for the imitation side to be included in the illustration. Video: -7s to -5s. Go cue onset: 0s. A) Healthy controls (N = 18), left hand imitation. B) Healthy controls (N = 18), right hand imitation. C) CBS patients (N = 11), left-hand imitation. D) CBS patients (N = 11), right-hand imitation. HC: Healthy controls; CBS: Corticobasal Syndrome.

The ERD associated with action observation was more pronounced in controls than CBS patients, for whom the ERD had a more posterior localization. Specifically, preand postcentral gyri as wells as the middle frontal gyrus, and parts of the inferior frontal gyrus, showed a weaker desynchronization bilaterally in patients (Fig. 4B; contralateral cluster: *tsum = 165.263, p = 0.041;* ipsilateral cluster: *tsum = 193.029, p = 0.034*). These areas served as regions of interest (ROI) in the following. The effect was specific to the beta band, covered the entire observation phase and outlasted it by about 300ms (Fig. 4C; contralateral cluster *tsum = 2496.215, p = 0.017;* ipsilateral cluster *tsum = 2479.829, p = 0.017*). Group differences in the alpha (8-12Hz) and gamma band (60-90Hz) were not significant.

**Figure 4.**
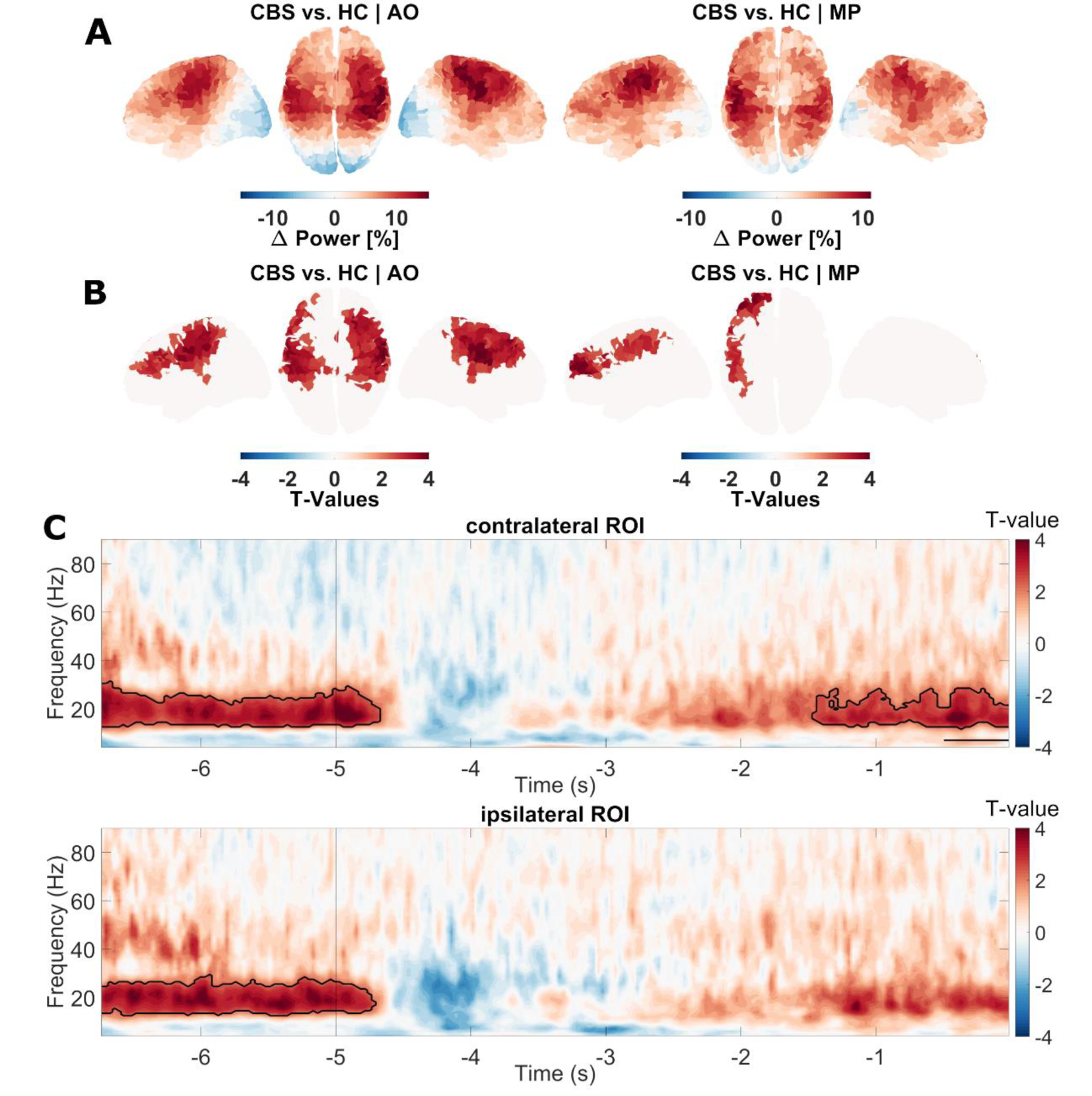
Comparison of event-related beta power desynchronization between CBS patients and healthy controls. A) Difference in group-average baseline-corrected beta-power (13-30Hz) for action observation (-6.75 s to -5 s) and movement preparation (-1.5s to 0s). Warmer colors indicate weaker event-related beta desynchronization (beta ERD) in patients. B) Whole-brain statistical comparison, beta ERD in CBS patients vs. HC. Non-significant changes masked. Highlighted areas in the left panel of B) served as regions of interest (ROI). C) Statistical comparison of time-frequency maps. Significant differences indicated by solid black contour. The black horizontal line marks the epoch containing movement in HC (see Fig. 2A). AO: Action observation; MP: Movement Preparation. CBS: Corticobasal Syndrome; HC: Healthy controls.

### Movement preparation

In the 5s following video offset, participants were waiting for the Go cue signaling imitation start. In this phase, we observed a second alpha/beta ERD, intensifying over time (Fig. 3, right panel). The pattern of desynchronization was more focal in comparison to action observation, with a clear peak in sensorimotor cortex contralateral to movement. This second ERD was again smaller in the CBS group, provided that trials were anchored to Go cue onset. Differences emerged in the pre-and postcentral gyri and middle frontal gyri contralateral to the response hand (Fig. 4B; cluster *tsum = 121.356, p = 0.039*). As for action observation, the difference within the ROI occurred in the beta band exclusively (Fig. 4C; contralateral cluster *tsum = 1471.227, p = 0.039; time range: -1.488s to 0s*). We note that part of the effect might have been mediated by differences in overt movement shortly before Go cue onset (Fig. 2A). When excluding the last 480ms of the trial, the statistical effect reduced to a trend (cluster *tsum = 90.431, p = 0.053*). No significant differences emerged in the alpha or in the gamma range.

### Trial Phase Comparison

Interestingly, the differences observed immediately before movement onset (-0.5s to 0s) vanished when trials were centered on movement onset rather than Go cue onset (Fig. 5; cluster *tsum = 8.87, p = 0.411*). In fact, the movement-locked ERD was remarkably similar for HC (Fig. 5A) and CBS (Fig. 5B) with respect to spatial extent, strength, and dynamics, indicating that movement initialization might not be altered in CBS patients.

**Figure 5.**
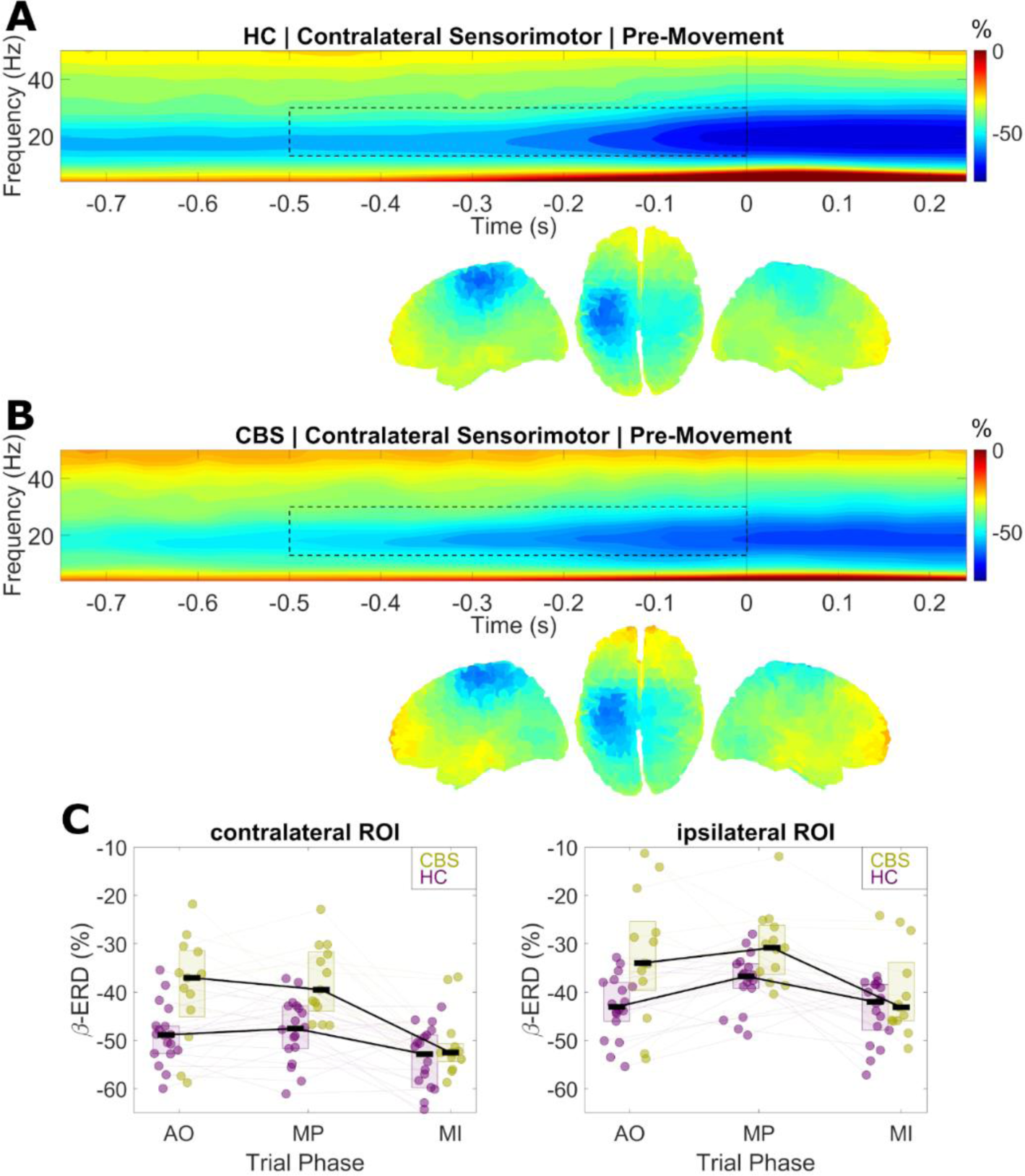
Beta power dynamics were similar for CBS patients and healthy controls immediately before movement onset. A) HC and B) CBS patients. Group-average time-frequency spectrum of sensorimotor cortex contralateral to imitation bodyside, anchored to movement onset (0s). Relative power change with respect to baseline (-7.75s to -7s) is color-coded. The dashed boxes indicate the time-frequency selection used in the source plots below. Leftand right-hand trials were pooled after mirroring the activity of left-hand trails across the sagittal plane (left hemisphere: contralateral to imitation). C) Beta ERD in the region of interest per trial phase. HC: Healthy controls; CBS: Corticobasal Syndrome; ROI: Region of interest; AO: Action observation; MP: Movement preparation; MI: Movement initialization. ERD: Event-related desynchronization.

In order to corroborate the absence of a group difference, we conducted a Bayesian analysis, assessing group effects on the beta ERD in three different trial phases: action observation (-7s to -5s relative to Go cue), movement preparation phase (-5s to 0s relative to Go cue) and movement initialization (-0.5s to 0s relative to movement onset). This analysis yielded strong evidence for an interaction between trial phase and group (*BF10 = 19.761*). Post-hoc comparisons yielded no evidence for a group difference during movement initialization (*BF10;contra.= 0.727; BF10;ipsi.= 0.697*), but strong evidence for action observation (*BF10;contra.= 29.399; BF10;ipsi.= 25.704*) and moderate evidence for movement preparation (*BF10;contra.= 8.751; BF10;ipsi.= 3.603*). The difference in effect size is illustrated in Fig. 5C.

### Go cue anticipation

The above analysis suggests that CBS patients and controls did not differ in motor cortical beta power suppression before moving (Fig. 5). Nevertheless, we observed group differences in beta ERD in the movement preparation phase (Fig. 4). We hypothesized that these findings can be reconciled by accounting for the difference in reaction time. CBS patients reacted slower than controls, which presumably went along with a slower suppression of beta power. In order to test this idea, we estimated the linear decay rate of beta power (beta slope) in the movement preparation phase and compared it across groups.

The estimated slopes of the beta ERD were smaller in the CBS group than in controls (Fig. 6A; contralateral ROI: *t(29) = 3.103, p = 0. 004;* ipsilateral ROI: *t(29) = 3.761, p < 0. 001*). Furthermore, the slope estimates of the contralateral hemisphere correlated with reaction times, confirming that steeper beta slopes are related to faster reaction times (Fig. 6B; Pearson partial correlation; contralateral ROI: *rBeta⨯RT|Group = 0.417, p = 0.022;* ipsilateral ROI: *rBeta⨯RT|Group = 0.212, p = 0.26*). The correlations remained significant when excluding the outlier with RT > 1s. These findings suggest that CBS patients were less oriented in time, resulting in insufficient suppression of beta power at Go cue onset and longer reaction times.

**Figure 6.**
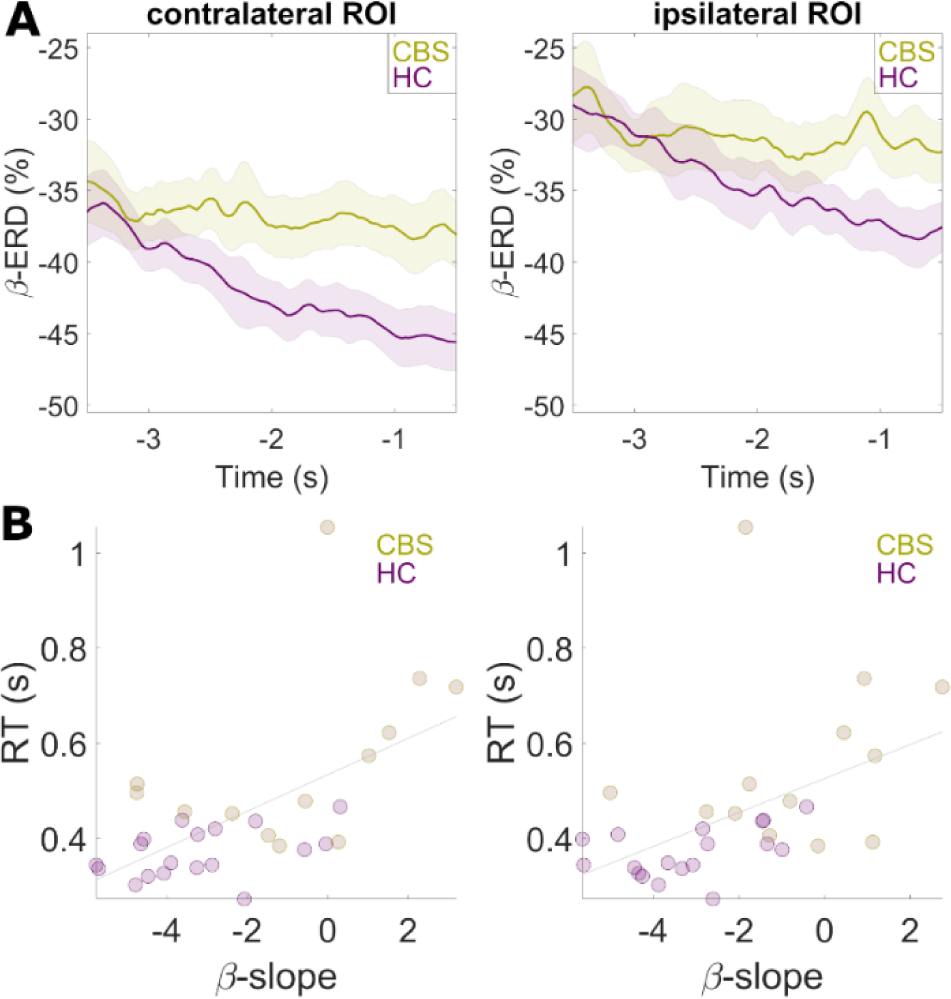
The rate of beta power suppression prior to the Go cue differed between CBS patients and healthy controls and correlated with reaction time. A. Beta power dynamics before Go cue presentation. The group means are displayed as colored lines, and the standard error is indicated by shaded areas. B. Linear decay rate of beta power (beta slope) vs. trial-median reaction time. Note that we omitted the last 480ms of the trial in this analysis because it contained hand movement (Fig. 2A). CBS: Corticobasal Syndrome; HC: Healthy controls; ROI: Region of interest; RT: Reaction time.

### Correlations between beta ERD and CBS symptoms

In CBS patients, we found no correlation (Spearman’s ρ) between the beta ERD in the contra- or ipsilateral region of interest and test scores quantifying apraxia (TULIA: *|ρ|< 0.375, p > 0.23*; Goldenberg: *|ρ| < 0.396, p > 0.182*), cognitive impairment (MoCA: *|ρ| < 0.385, p > 0.194*), or parkinsonism (UPDRS III: *|ρ| < 0.251, p > 0.409*). We neither found correlations between test scores and beta slope estimates (*|ρ| < 0.509, p > 0.076*).

## Discussion

### Summary

Little is known about the patho-electrophysiolgy of CBS. In this paper, we provide a comprehensive characterization of the electrophysiological differences between CBS patients and healthy age-matched controls in an observe-to-imitate task, requiring functions believed to be impaired in CBS, such as visuo-motor mapping and motor preparation. We found that action observation and GO cue anticipation were associated with beta power desynchronization in motor and parietal areas. These modulations, timed to action-relevant, visual information presented before movement onset, were weaker in CBS patients than in controls. The degree of beta power suppression immediately before movement onset, in contrast, was not different between patients and controls, suggesting that the effects observed here are neural correlates of selective deficits in visuo-motor mapping and implicit learning of temporal structure, respectively, rather than of impaired movement initialization.

### Action Observation

Functional magnetic resonance studies revealed a comprehensive brain network encompassing both subcortical and cortical regions engaged in the observation of meaningful actions. Subcortically, activation occurs in the cerebellum (Gazzola and Keysers 2009; Molenberghs et al. 2012), the basal ganglia (Errante and Fogassi 2020; Errante et al. 2023), and the thalamus (Errante and Fogassi 2020; Errante et al. 2023). Cortically, frontal regions such as premotor and precentral cortex are involved, alongside the supplementary motor area, primary somatosensory cortex, parietal and occipital cortex (Caspers et al. 2010; Molenberghs et al. 2012; Hardwick et al. 2018; Errante and Fogassi 2020; Errante et al. 2023).

On the electrophysiological level, action observation is associated with a suppression of beta oscillations in sensorimotor areas (Cochin et al. 1998; Hari et al. 1998; Babiloni et al. 2002; Muthukumaraswamy and Johnson 2004; Caetano et al. 2007; Sebastiani et al. 2014; Pavlidou et al. 2014b, 2014a; Kilner et al. 2009). In agreement with the current study, a previous study localized this beta desynchronization to frontoparietal areas, and primary sensorimotor areas in particular (Sebastiani et al. 2014). Concerning the lateralization of beta desynchronization during action observation there is conflicting evidence. One study reported bilateral desynchronization (Babiloni et al. 2002) while another study reports contralateral desynchronization with respect to the target stimulus (Kilner et al. 2009). We found that beta desynchronization during action observation shows a mild contralateral predominance, which might have resulted from the need to imitate the observed action with one hand later in the trial.

Providing movement context is known to affect the beta desynchronization associated with action observation, although the nature of these effects is not fully understood. Muthukumaraswamy et al. (2004), for example, found that the observation of meaningless movements leads to less beta suppression than the observation of goal-directed movements. Pavlidou et al. (2014b), in contrast, found that biologically implausible movements rather than plausible movements are related to larger beta power decreases, which was attributed to a difference in effort when matching visual information onto motor representations. These reports imply that beta modulations emerging during action observation are related to cognitive processes rather than kinematics only.

### Similarities between action observation, motor imagery, and movement execution

The electrophysiological signature of action observation is remarkably similar to that of movement execution and motor imagery. Before participants begin to move (Toro et al. 1994; Fairhall et al. 2007), or imagine a movement (Pfurtscheller and Neuper 1997; Schnitzler et al. 1997; McFarland et al. 2000; Eaves et al. 2016), beta power decreases in primary sensorimotor areas. Given the substantial evidence supporting an inhibitory role of beta oscillations in motor control, stemming from studies on Parkinson’s disease (Kühn et al. 2004; Swann et al. 2011; Alegre et al. 2013; Toledo et al. 2014) and response inhibition (Swann et al. 2012; Picazio et al. 2014; Schaum et al. 2021), this beta power suppression likely reflects a transient disinhibition of primary sensorimotor cortex, which is otherwise constantly inhibited by the response inhibition network, including pre-supplementary motor area (Swann et al. 2012; Picazio et al. 2014; Schaum et al. 2021), inferior frontal cortex (Swann et al. 2012; Picazio et al. 2014; Schaum et al. 2021) and the subthalamic nucleus (Kühn et al. 2004; Alegre et al. 2013; Chen et al. 2020). Notably, this transient disinhibition does not necessarily result in overt movement. It rather reflects an “active network state” (Pogosyan et al. 2009; Little et al. 2019; Muralidharan and Aron 2021) common to action observation, motor imagery and movement execution. In line with this interpretation, a recent meta-analysis of fMRI studies has demonstrated that the activation maps of action observation, motor imagery, and motor execution considerably overlap in premotor, sensorimotor, and rostral parietal areas (Hardwick et al. 2018).

### Differences between action observation, motor imagery and movement execution

Besides the abovementioned similarities, several differences have been identified between action observation, motor imagery, and motor execution. On the electrophysiological level, these differences appear to be rather subtle. In one study, observation of a rhythmic action and concurrent motor imagery of a different rhythmic action induced a stronger beta ERD than either action observation or motor imagery alone (Eaves et al. 2016). When the rhythm of the imagined action needed to be synchronized to the observed action, the authors observed an additional prefrontal beta ERD, presumably indicating increasing demands in cognitive control (Eaves et al. 2016). Additionally, there is evidence that motor imagery, but not action observation, modulates corticospinal excitability (Meers et al. 2020). Both of these studies point towards partial independence between action observation and motor imagery.

Similarly, imaging studies support partial independence between action observation, motor imagery and motor execution by revealing spatial differences in the associated brain activity patterns. Notably, movement execution engages a rather focal cortical network centered on primary sensorimotor areas, with a limited activation of premotor and inferior parietal regions (Hardwick et al. 2018). Action observation and motor imagery, in contrast, recruit a more extended network, including premotor, pre-SMA and various parietal areas (Hardwick et al. 2018). These additional frontoparietal regions might be required for visuo-motor mapping, i.e. the integration of the visual percept and the motor representation of an action, which is particularly important in observe-to-imitate tasks.

Substantial parts of the frontoparietal network are affected by pathological changes in CBS, likely explaining why we observed the strongest CBS-related alterations in the action observation phase. Pathological alterations include both gray matter degeneration (Huey et al. 2009; Josephs et al. 2010; Whitwell et al. 2010; Dutt et al. 2016; Matsuda et al. 2020) and damage to white matter tracts that link frontal with parietal cortices and/or subcortical regions, such as the superior longitudinal fasciculus (Ferrea et al. 2022; Uchida et al. 2023). The subcomponents of the superior longitudinal fasciculus linking parietal and frontal regions (Makris et al. 2005; Nakajima et al. 2020), in particular, might facilitate visuo-motor mapping during action observation (Hecht et al. 2013).

### Movement preparation

The disinhibition of motor cortex during action observation, reflected by the first beta ERD in our task, was likely a direct consequence of action observation, and thus largely independent of learning. The ERD timed to the Go cue, in contrast, was presumably contingent on learning the trial’s temporal structure, including the constant interval between video offset and Go cue onset. Previous literature has demonstrated that beta power adapts to the timing of a predictable, upcoming target stimulus (van Ede et al. 2011; Heideman et al. 2018). In line with these studies, Tzagarakis et al (2010) demonstrated that beta desynchronization is modulated by response uncertainty. Thus, the reduced pre-Go beta modulation in the CBS cohort is likely the result of uncertainty regarding the onset of the Go stimulus. The fact that patients have a slower ERD might thus be indicative of an impairment in learning temporal structure. In line with this idea, beta desynchronization correlated with reaction time, confirming previous reports (Perfetti et al. 2011; Tzagarakis et al. 2010). The deficit in learning temporal structure could potentially be due to widespread neurodegeneration in frontal cortex, parietal cortex and basal ganglia, which are known to be involved in time estimation (Coull et al. 2011; Coull et al. 2013). Premotor and parietal cortex are particularly relevant for anticipation of cues and response preparation (Coull et al. 2011).

### Movement initialization

Previous findings in humans and non-human primates suggest that beta power must be suppressed below a certain threshold for movement initialization (Heinrichs-Graham and Wilson 2016; Khanna and Carmena 2017). Here, we observed a slower beta power suppression and prolonged reaction times in CBS patients relative to controls, but no difference with respect to the level of beta power suppression at movement onset. This finding suggests that the power threshold for movement initialization, relative to baseline, is similar in both groups. The temporal timing of beta power suppression to the task, however, might be pathologically altered in CBS patients.

### Clinical implications

Our results evidence pathological alterations of beta desynchronization in CBS patients, presumably caused by neurodegeneration in brain circuits involved in visuo-motor mapping and implicit learning of temporal structure. These electrophysiological alterations can be evaluated easily by presenting a tool-use video, coupled with the instruction to imitate, while recording MEG/EEG. Unlike motor imagery, this task does not require a high level of patient compliance. These properties make our approach potentially interesting for translation into diagnostic tools, that might be useful for differentiating Parkinson syndromes in the future.

### Limitations

We did not find significant correlations between clinical scores and beta power desynchronization. This might be due to the relatively small sample size (N=13) and the symptomatic variability in our patient cohort. Alternatively, it is conceivable that the observed alterations of brain activity might relate more to neurodegeneration per se than to clinical symptoms, which are known to have different long-term dynamics in neurodegenerative diseases (Armstrong et al. 2013; Aiba et al. 2023). Lastly, our study lacked several experimental conditions of interest, such as a motor imagery task, action observation without the need to imitate or action observation from different perspectives.

## Conclusion

The processing of observed actions is pathologically altered in CBS, likely reflecting a selective deficit in visuo-motor mapping. In addition, CBS patients show suboptimal timing of beta suppression to the task, presumably due to deficits in implicit learning of temporal structure.

## Acknowledgement

This study was funded by Else Kröner Fresenius Stiftung and Brunhilde Moll Stiftung.

## Data Availability

We do not have permission to share the data publicly.

