## Supplementary Material for "Altered Cortical Network Dynamics during Observing and Preparing Action in Patients with Corticobasal Syndrome"

Moorenstraße 5

40225 Düsseldorf, Germany

**Word count:** 5077

**Running title**: Observation and preparation of action in patients with Corticobasal Syndrome

**Financial disclosure / Conflict of interest:** Nothing to report.

**Founding:** This study was funded by the Else Kröner Fresenius Stiftung and Brunhilde Moll Stiftung.

### Supplementary Materials

**Table 1: Number of trials per block after cleaning.**

| Participant | RH01 | LH01 | RH02 | LH02 | Total |
|---|---|---|---|---|---|
| **CBS01** | 29 | 32 | 34 | 27 | 122 |
| **CBS02** | 37 | 22 | - | - | 59 |
| **CBS03** | 27 | 18 | - | - | 45 |
| **CBS08** | 30 | 28 | - | 29 | 87 |
| **CBS10** | - | 28 | - | - | 28 |
| **CBS11** | 27 | 27 | 24 | 31 | 109 |
| **CBS12** | 10 | 26 | | 26 | 62 |
| **CBS15** | 28 | 33 | 33 | 32 | 126 |
| **CBS16** | 27 | 33 | 34 | 30 | 124 |
| **CBS18** | 17 | 29 | 15 | 24 | 85 |
| **CBS19** | 34 | 29 | 26 | 0 | 89 |
| **CBS20** | 36 | 36 | 34 | 26 | 132 |
| **CBS21** | 27 | 29 | - | - | 56 |
| **HC11** | 33 | 31 | 32 | 32 | 128 |
| **HC12** | 34 | 34 | 30 | 28 | 126 |
| **HC13** | 32 | 31 | 34 | 35 | 132 |
| **HC14** | 27 | 28 | 29 | 27 | 111 |
| **HC15** | 33 | 23 | 30 | 32 | 118 |
| **HC16** | 39 | 38 | 31 | 36 | 144 |
| **HC17** | 32 | 31 | 36 | 37 | 136 |
| **HC18** | 30 | 24 | 26 | 21 | 101 |
| **HC24** | 18 | 15 | 28 | 26 | 87 |
| **HC25** | 26 | 36 | 32 | 34 | 128 |
| **HC26** | 20 | 26 | 19 | 27 | 92 |
| **HC29** | 20 | 21 | 28 | 24 | 93 |
| **HC30** | 31 | 35 | 35 | 31 | 132 |
| **HC31** | 32 | 34 | 26 | 30 | 122 |
| **HC32** | 33 | 30 | 32 | 31 | 126 |
| **HC33** | 40 | 40 | 39 | 40 | 159 |
| **HC34** | 37 | 37 | 34 | 39 | 147 |
| **HC35** | 39 | 39 | 40 | 39 | 157 |

CBS: Corticobasal Syndrome, HC: Healthy controls.
